# Using Citizen Science to build baseline data on tropical tree phenology

**DOI:** 10.1101/2020.08.28.271155

**Authors:** Geetha Ramaswami, Swati Sidhu, Suhel Quader

## Abstract

Large-scale and long-term understanding of the phenology of widespread tree species is lacking in the tropics, and particularly in the Indian subcontinent. In the absence of baseline information, the impacts of climate on tree phenology, and thus on trophic interactions downstream of tree phenology, are also poorly understood. Citizen scientists can help bridge this gap by contributing simple, technology-based information over large spatial scales and over the long term. In this study, we describe an India-wide citizen science initiative called SeasonWatch, with preliminary insights into contributor behaviour and species phenology. Over a period of 8 years, between 2011 and 2019, cumulative contributor numbers have increased every year, although consistent contribution remains constant and low. The phenological patterns in the 4 most-observed species (Jackfruit *Artocarpus heterophyllus* Lam., Mango *Mangifera indica* L., Tamarind *Tamarindus indica* L., and Indian Laburnum *Cassia fistula* L.) are described, with discernible seasonal peaks in flowering and fruiting. Seasonal peaks are influenced by tree phenology reported in the south Indian state of Kerala, which has the maximum number of contributors and most number of observations per contributor, comprising 89% of all observations. We look in detail at the flowering phenology of one particular species, *Cassia fistula*, which appears to show aberrant phenology, reflecting a potential shift away from historical baselines. Latitudinal patterns in the phenology of widespread species such as *Mangifera indica* are also discernible from 4 seasonal bioblitz events organised during 2018-19, with trees in lower latitudes exhibiting flowering and fruiting phenology earlier than the higher latitudes. We conclude that there are signs of shifts in phenological patterns, as in the case of *C. fistula*, and extend a call for action to sustain long-term interest and participation by contributors to develop a baseline for common tropical tree species that can be used to understand long-term consequences of climate change on tropical tree phenology.

## Introduction

Cyclic patterns of growth and reproduction - or phenology - of living organisms are seasonal and highly sensitive to the environment (Sparks and Carey, 1995). Discernible changes outside of the known variability in the phenology of organisms are often indicative of underlying changes in large-scale climate patterns. In temperate regions, higher temperatures are related to the onset of spring phenophases, such as flowering and leaf unfolding (Menzel et al 2006). Increasingly, dates of bud-burst have been demonstrated to advance when compared with long-term averages due to advancement in the warmer season, increase in winter and spring temperatures, and even effects of urbanisation such as pavements and light at night-time (e.g. Menzel et al 2006, Fu et al 2012, Chen et al 2016, ffrench-Constant et al 2016). In the tropics, recent long-term phenological studies have shown that the onset, persistence and frequency of reproductive phenophases are affected by solar irradiance (Babweteera et al. 2018; Chapman et al. 2018; Wright and Calderón 2018). In other tropical systems, reproductive phenology has been reported to be affected by precipitation; fruiting intensity often increases with higher rainfall (Dunham et al. 2018; Mendoza et al. 2018). However, the impacts of changes in baseline environmental conditions on tropical tree phenology are poorly understood.

The bulk of our understanding of phenological patterns in plants, and their response to climatic change, comes from Europe and North America (Bertin 2008). Apart from contemporary studies, this understanding is supplemented by a number of historical data sets collected by hobbyists and naturalists, that serve as baselines for comparing current phenological patterns (the Marsham phenological record, see Sparks and Carey, 1995). Some of the largest historical datasets on phenology are from temperate regions, including the Kyoto cherry blossom dataset which is over 1300 years old (Aono and Kazui, 2008). Most contemporary phenology monitoring networks are also situated in temperate latitudes (Cleland 2007). Information on tropical phenology is lacking both in contemporary as well as historical studies, and large-scale or generalizable responses to climate are not known (Kushwaha and Singh, 2008).

Arriving at generalizable trends for phenological responses to climate requires data collected over large spatial scales and over multiple years, and integrating different datasets (Morisette et al 2008). Intensive research efforts at such scales are often not possible due to resource and personnel limitations. Large-scale, long-term observations, however, can be facilitated by citizen scientists (Schwartz, Betancourt, and Weltzin, 2012). Citizen science involves members of the public in scientific studies to collectively generate information around issues of larger public concern. The term, initially coined by the Cornell Lab of Ornithology in 1995 (Bonney 2016), has since been used to include a multitude of projects such as mapping and documenting biodiversity (iNaturalist, Project Noah, PlantWatch, eBird), population dynamics (StreamWatch), bird migration (North American Bird Phenology Program), and plant phenology (Nature’s Calendar, SeasonWatch). Citizen science projects are designed such that interested people, with or without a formal training in science, volunteer their time and collect information required to build and co-create datasets of interest (Dickinson et al 2012, Wiggins and Crowston 2011).

Worldwide, citizen science initiatives have contributed to large-scale understanding of phenology, including the quantification of climate change impacts on plant phenology. For instance, under different climate change scenarios, the flowering and fruiting phenology of *Vaccinium membranaceum* is predicted to advance on average by 35 and 36 days respectively in the USA. These predictions are based on the current understanding of the environmental correlates of reproductive phenology of *V. membranaceum* from data contributed to the citizen science initiative USA National Phenology Network along with other long-term data sources (Prevey et al 2020).

Within India, multiple past or ongoing efforts have gathered long-term phenology data in different sites that is contextualized for the region and for the species occurring therein (Ramaswami et al 2019). Among them, SeasonWatch, a citizen science initiative that started in 2010, is collating data on phenology of common Indian trees at a country-wide level, with the ultimate aim of understanding geographical variation and the effects of climate change on the phenology of widespread species. Apart from the formal literature (floras, scientific articles, tree guides etc.), there is also anecdotal and cultural information on the expected phenology of trees. For instance, the festival of Vishu (typically in the 15th week of the year, between 13 and 15 April) in Kerala is associated with flowering in *Cassia fistula*, with inflorescences forming an integral part of the festivities. In recent years, there have been anecdotal and media reports in Kerala, of the flowers of *C. fistula* being unavailable during the festival of Vishu, as well as aberrant flowering during other times of the year.

Here, we describe SeasonWatch as an attempt to create a baseline dataset on tropical tree phenology using citizen science. We illustrate this by exploring phenological patterns of selected tree species across space and time. We also summarise contributor history and consistency (details in Methods section), regional representation of trees, and assess whether SeasonWatch data can be used to understand probable shifts in phenological patterns of trees in the future. Specifically, we provide summaries of 1) contributor behaviour in the context of data upload patterns, 2) tree behaviour and phenological patterns over time, 3) flowering phenology of *C. fistula* over time, 4) phenological patterns over space using data for *Mangifera indica*. We conclude with potential research questions of interest in the future.

## Methods

### Project summary

We summarise data contributed to SeasonWatch over a period of six years between 1 January 2014 and 31 December 2019. Data contribution requires registration with the SeasonWatch project. Registered contributors upload information on the phenology of 136 common, widespread (or locally abundant), native and non-native (planted along avenues and roadways, or invasive species of concern) woody plant species (list available at www.seasonwatch.in/species.php), via the project website (www.seasonwatch.in) or a freely downloadable Android mobile phone app.

### Contributor summary

SeasonWatch has two main types of contributors: schools (teachers and students, typically of grades 6-9) and individuals (nature enthusiasts; students at universities). School teachers learn about participating in the project through state and district-level teacher-training workshops, or through workshops at schools, and implement the project in their school premises with groups of students. SeasonWatch typically partners with a regional organisation in order to facilitate local outreach and training.

Contributors can choose to make an observation on a tree only once (‘Casual’ henceforth) or register a tree with the project and make observations on its phenology every week (‘Regular’ henceforth). A large proportion of the Regular data is contributed by schools and these data are consistently received throughout the school year. Regular observations allow for understanding individual-level variation in seasonal phenology of trees, keeping location constant. Casual observations have been collected largely through four ‘bioblitz’ events held in December 2018, and in March, June, and September 2019. The aim of the 4-day ‘bioblitz’ events was to create a snapshot of phenology across India in different seasons. Casual observations allow for understanding latitudinal variation in species phenology.

### Observations summary

Each contributed data point contains the following information - tree location, date and time of observation, date of upload, species identity, and an estimate of leaf, flower and fruit quantity on the observed tree. ‘Leaf’, ‘flower’ and ‘fruit’ are further categorised into finer phenological stages (Table 1). The quantity of each phenophase is noted as one of the following - ‘None’, ‘Few’ or ‘Many’. ‘None’ corresponds to the absence of the phenophase. If a phenophase is observed on 1/3rd or less of the tree canopy, it is marked as ‘Few’ by a contributor, indicating a lower volume of the tree having the reported phenophase. If a phenophase is observed on more than 1/3rd of the tree canopy, it is marked as ‘Many’ by a contributor, indicating a larger volume of the tree having the reported phenophase. ‘Many’ is considered as the phenophase peak at the level of each individual tree. On any given day, only one of these categories can be reported for a phenophase. Along with phenology observations, contributors can optionally note the presence of herbivores, pollinators and dispersers, and also make additional notes on natural history, interactions of interest etc. All phenology data contributed to SeasonWatch are available to any contributor or non-contributor, upon request.

**Table 1:** Tree phenophases recorded by citizen scientists in SeasonWatch. Phenophases are recognised based on descriptors

| Phenological indicator | Phenophase | Descriptors |
| --- | --- | --- |
| Leaf | Fresh | Leaf bud; recently emerged; small-sized; light green or pigmented |
|  | Mature | Full-sized; dark green |
|  | Dying | Necrotic; discoloured; extreme herbivory |
| Flower | Bud | Floral bud |
|  | Male* | Androecium only, corolla unfurled |
|  | Female* | Gynoecium only, corolla unfurled |
|  | Open | Bisexual flowers, corolla unfurled |
| Fruits | Unripe | Small-sized; dark green |
|  | Ripe | Full-sized; mature fruit coloration |
|  | Open* | Fruit opened, seeds dispersed |

\*Phenophase present only in a few species. ‘Dying’ Leaf and ‘Open’ fruit phenophases were introduced in the year 2018 in order to capture deciduousness and seasonal splitting of pods/capsules respectively of certain species.

Based on these contributed observations, we quantify the following:

#### 1) Contributor behavior and data upload patterns

We summarise contributor behaviour in the form of overall participation (in making Regular and Casual observations) from each Indian state, and for each year. We quantify contributor consistency as a measure of continuity in Regular observations on registered trees. A contributor is defined as ‘consistent’ if she makes phenology observations in 23 or more unique weeks in a year.

#### 2) Tree behaviour and phenological patterns over time and space

We summarise average weekly phenological responses of tree species over a period of 6 years. We assessed Regular phenology data for patterns of phenology using data contributed between 1 January 2014 and 31 December 2019. Since contributors are encouraged to record phenology once a week, we also consider the week as the smallest unit of time to report average patterns across species. Tree behaviour was quantified as the average proportion of trees displaying a particular phenophase in a week.

#### 3) Flowering in *Cassia fistula*

We assessed long-term patterns of flowering in *C. fistula*, by combining Regular and Casual data for this species between 2014 and 2019. Tree behaviour was measured as the proportion of trees displaying the floral phenophase ‘open flower’, comparing seasonal patterns in any flowering (i.e., open flowers stages reported as ‘few’ or ‘many’) with those in full bloom (i.e., ‘open flowers’ reported as ‘many’). The denominator in these proportions is the total number of individuals for which phenology was reported in each week. We visually compare peaks in the proportion of trees showing any flowering with those that were reported as being in full bloom to understand whether overall flowering patterns in *C. fistula* are different from that which is anecdotally expected.

#### 4) Patterns over space

Country-wide spatial phenological patterns in tree phenology were based only on Casual observations during the four bioblitz events. We plotted the spatial locations of phenophases of interest in the most observed species on a map to understand latitudinal variation in species phenology.

## Results

We summarise the overall and regional patterns of contributors and species phenology in SeasonWatch. Since 2011, 1050 individuals together with students from 889 schools from across India have contributed 351,481 phenology observations. Of these, 283,940 are repeat observations on registered trees, while 67,541 are one-time observations contributed during quarterly bioblitz events in 2018/19.

### 1) Contributor behaviour and data upload patterns

The total number of unique contributors with at least one valid phenology observation has been variable since 2011, with an increase in both individual and schools since 2017 (Fig 1a). The number of consistent contributors has remained constant and low, since 2011, even though there is an increase in total number of contributors (Fig 1b). The south Indian state of Kerala has the largest number of contributors as well as the largest number of observations per contributor (Fig 1c).

**Fig 1:**
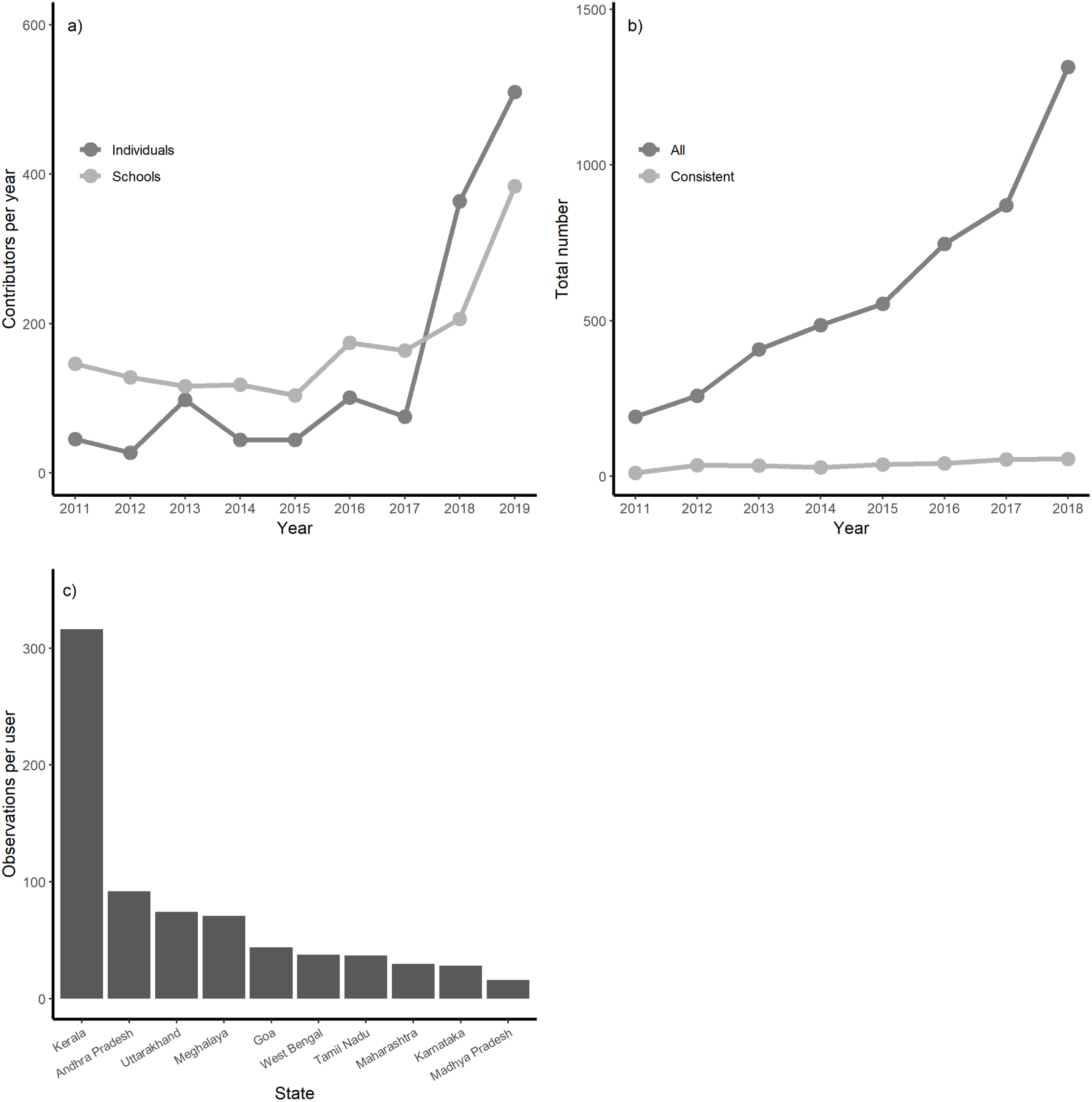
Contributor behaviour over time. a) number of schools and individuals with at least one valid phenology observation between 1 January 2014 and 31 December 2019. b) Number of contributors added per year and number of consistent contributors with at least one observation made in at least 23 weeks of the year. c) per-contributor average number of observations from different states of India

### 2) Tree behaviour and phenological patterns over time

The species with the most number of observations differed between the two types of observation. 62 species are represented by 100 or more Regular observations and 73 are represented by 100 or more Casual observations. Five species with the most number of Regular observations are *Artocarpus heterophyllus* Lam. (Jackfruit), *Mangifera indica* L. (Mango), *Tamarindus indica* L. (Tamarind), *Phyllanthus emblica* L. (Indian gooseberry), and *Cassia fistula* L. (Indian Laburnum, Figure 2a). Five species with the most number of one-time observations are slightly different: *Tectona grandis* (Teak), *Cocos nucifera* L. (Coconut palm), *M. indica, A. heterophyllus* and *Azadirachta indica* A.Juss. (Neem, Figure 2b).

**Fig 2:**
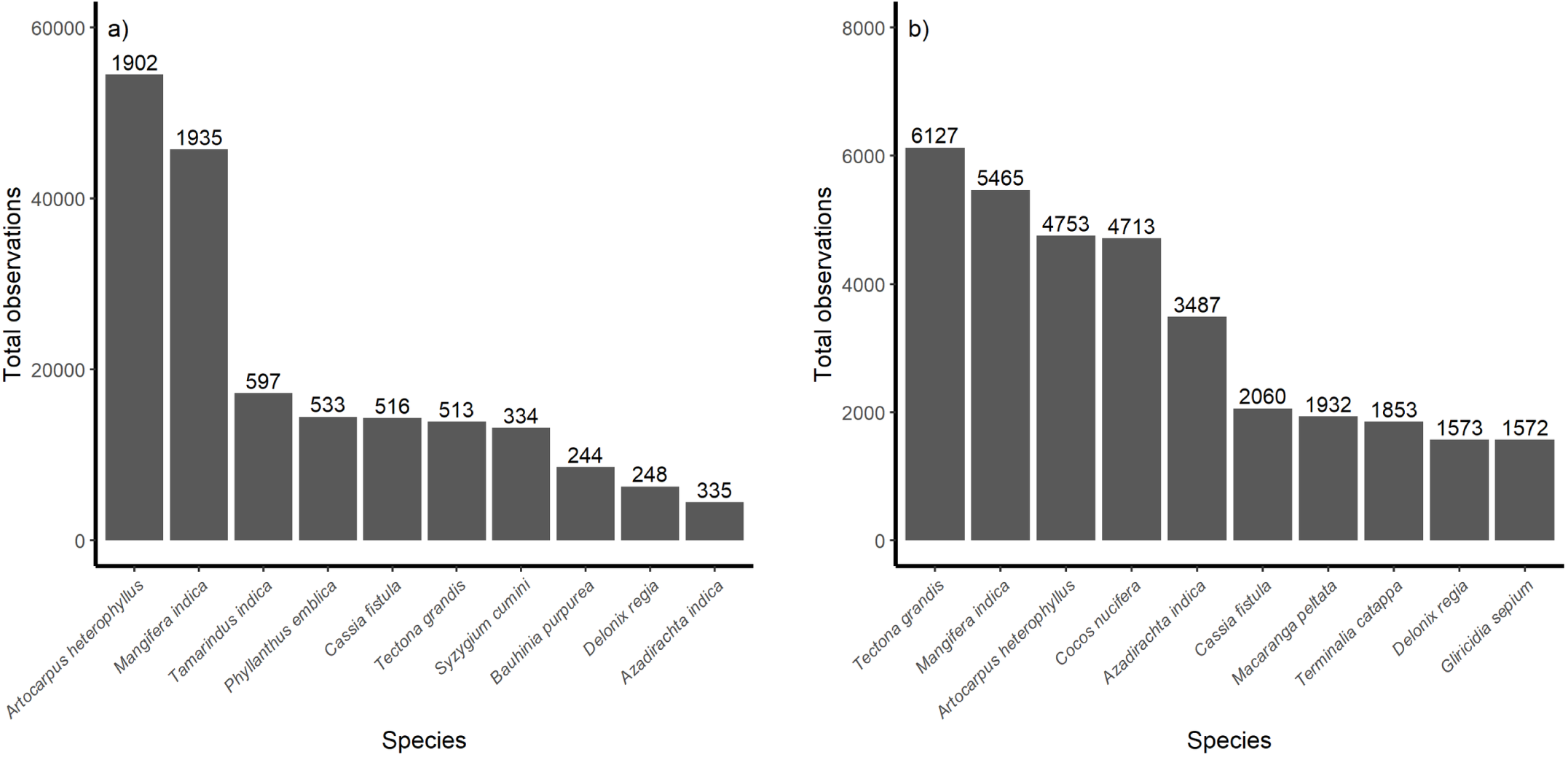
Total number of observations per species from 1 January 2014 to 31 December 2019. a) ‘Regular’ observations are repeat observations on registered trees; numbers on top of the bars indicate the number of registered individual trees for that species across India. b) ‘Casual’ observations are made once on any individual tree; number of observations is thus, the same as the number of individual trees observed, assuming that every casual observation is made on a different tree

The seasonal patterns in the appearance of leaf, flower and fruit based on Regular observations are given in Figure 3(a-d) for the 4 species with the most number of repeated observations. Peaks in flowering and fruiting are clearly discernible in *M. indica, A. heterophyllus* and *T. indica*, but not as apparent in *P. emblica*. The majority of Regular observations are contributed from the south Indian state of Kerala, and thus the peaks in phenology are most reflective of the seasonal response of trees at these latitudes (8.3218 - 12.7890 degrees N). Sample sizes for these patterns are provided in Appendix 1.

**Figure 3:**
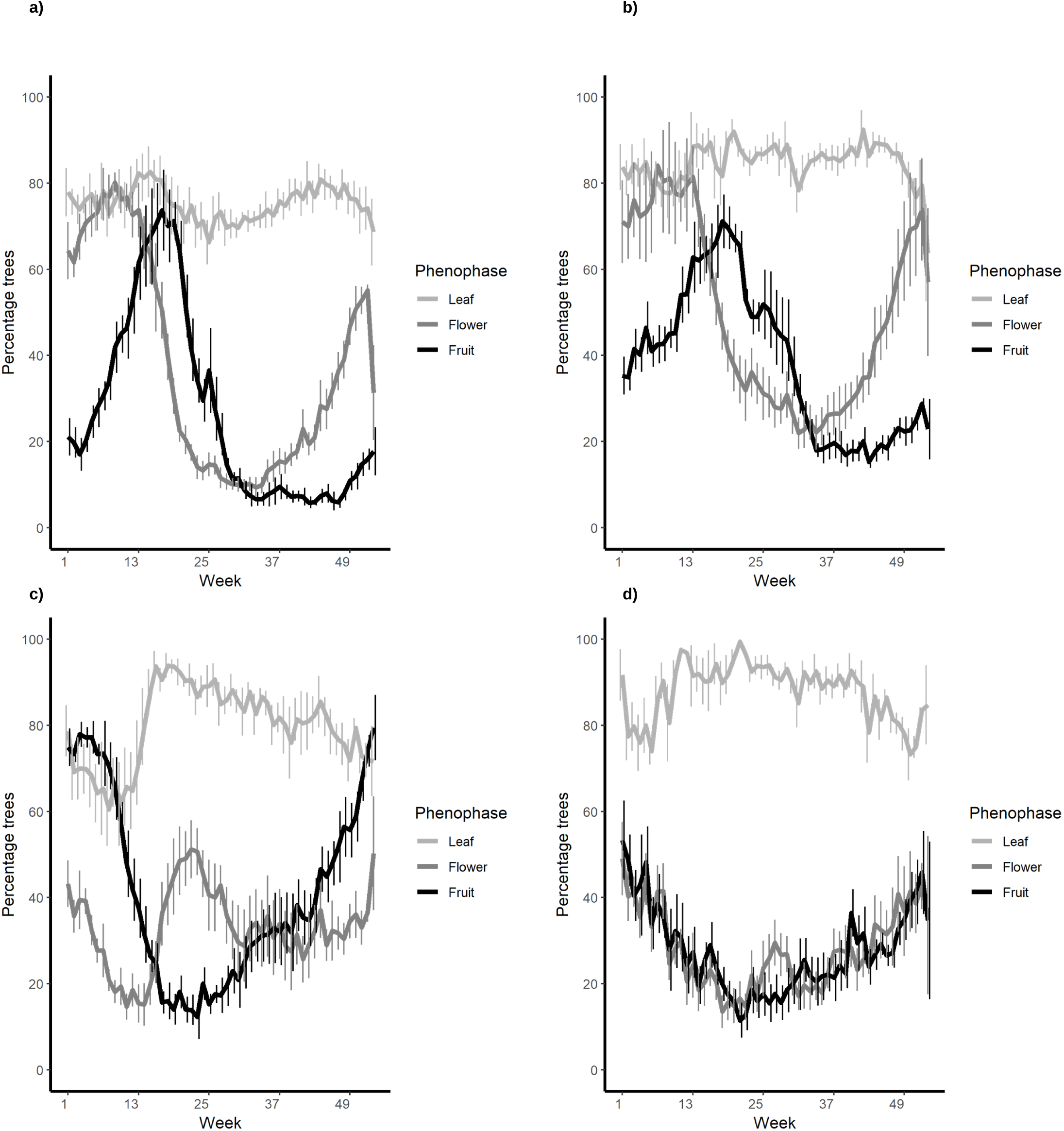
Phenology of four most observed species across India. a) *Mangifera indica*, b) *Artocarpus heterophyllus* and c) *Tamarindus indicus* show clear seasonal peaks in flowering and fruiting, while d) *Phyllanthus emblica* shows moderate, persistent flowering and fruiting through the year. Bars indicate standard error across 5 years, sample sizes for these patterns are given in Appendix 1.

### 3) Flowering phenology of C. fistula

Between 2014 and 2019, we found that up to 40% of trees (on average) of this species were reported to have open flowers in nearly every week of the year. However, nearly 70% of observed trees on average, were reported to be in full bloom in the 13th week of the year, indicating a flowering peak. This declined to 50% of observed trees on average by week 15 (Figure 4). This could be an indication of a shift in flowering from the expected time of week 15 in the year. Sample sizes for these observations, however, are low and highly variable (weekly ranging from 17 to 124 trees) and more observations are required in order to determine whether there is indeed a shift from the expected time of flowering in this species.

**Figure 4:**
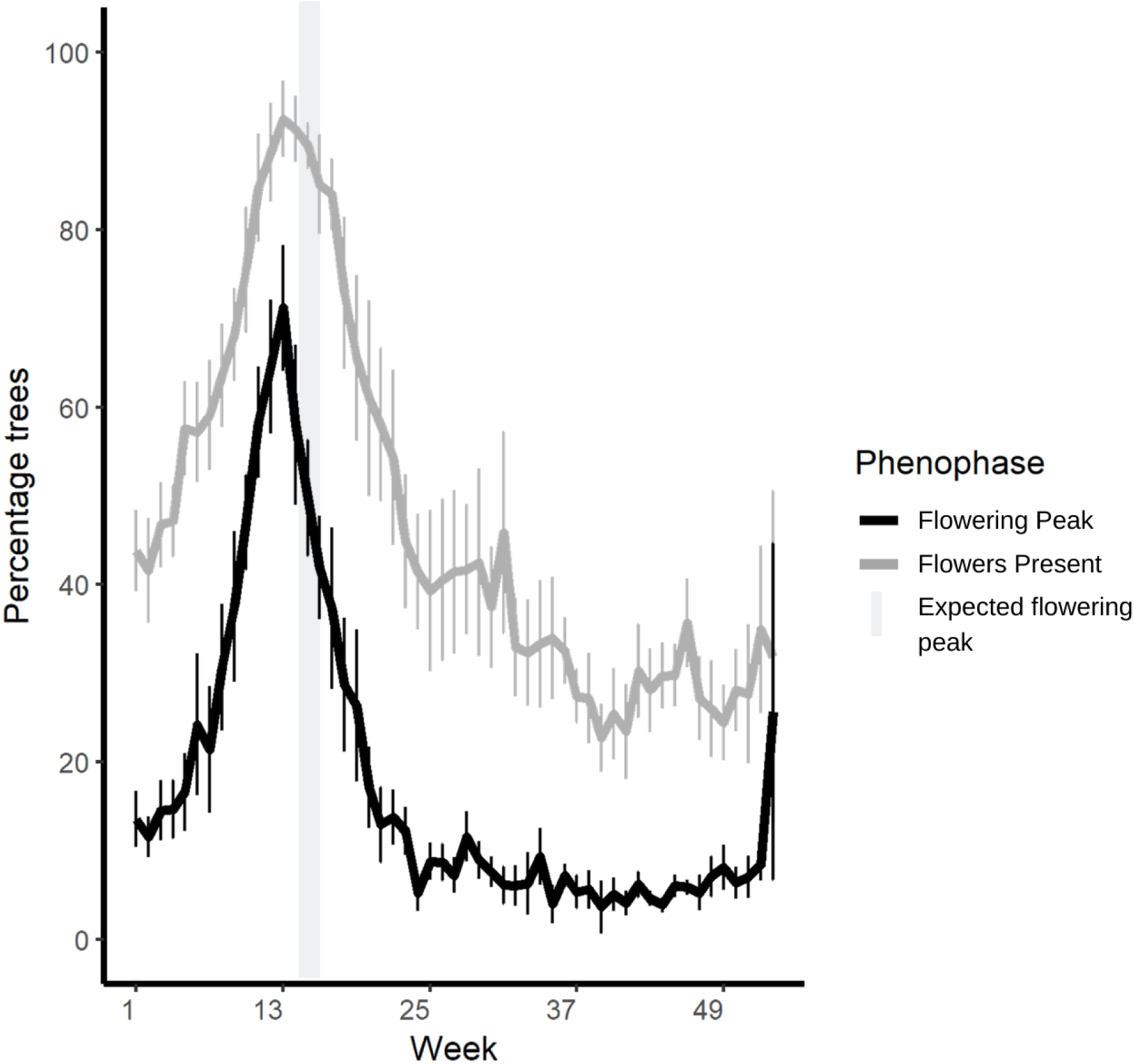
Proportion trees with phenophase indicated as ‘present’ or ‘many’ (indicating a perception of phenophase peak by the contributor). Flowering in Cassia fistula compared against the expected peak flowering time indicated by the cultural festival of Vishu in Kerala.

### 4) Phenological patterns over space

The number of contributors and observations during 4 bioblitz events was variable, with the maximum participation in December 2018 and, maximum observations contributed in June 2019 by fewer contributors (Fig 5a and b). The most reported species from different latitudes varied across the four bioblitz events and latitudinal patterns could be assessed for only the most widespread, commonly seen tree species.

**Figure 5:**
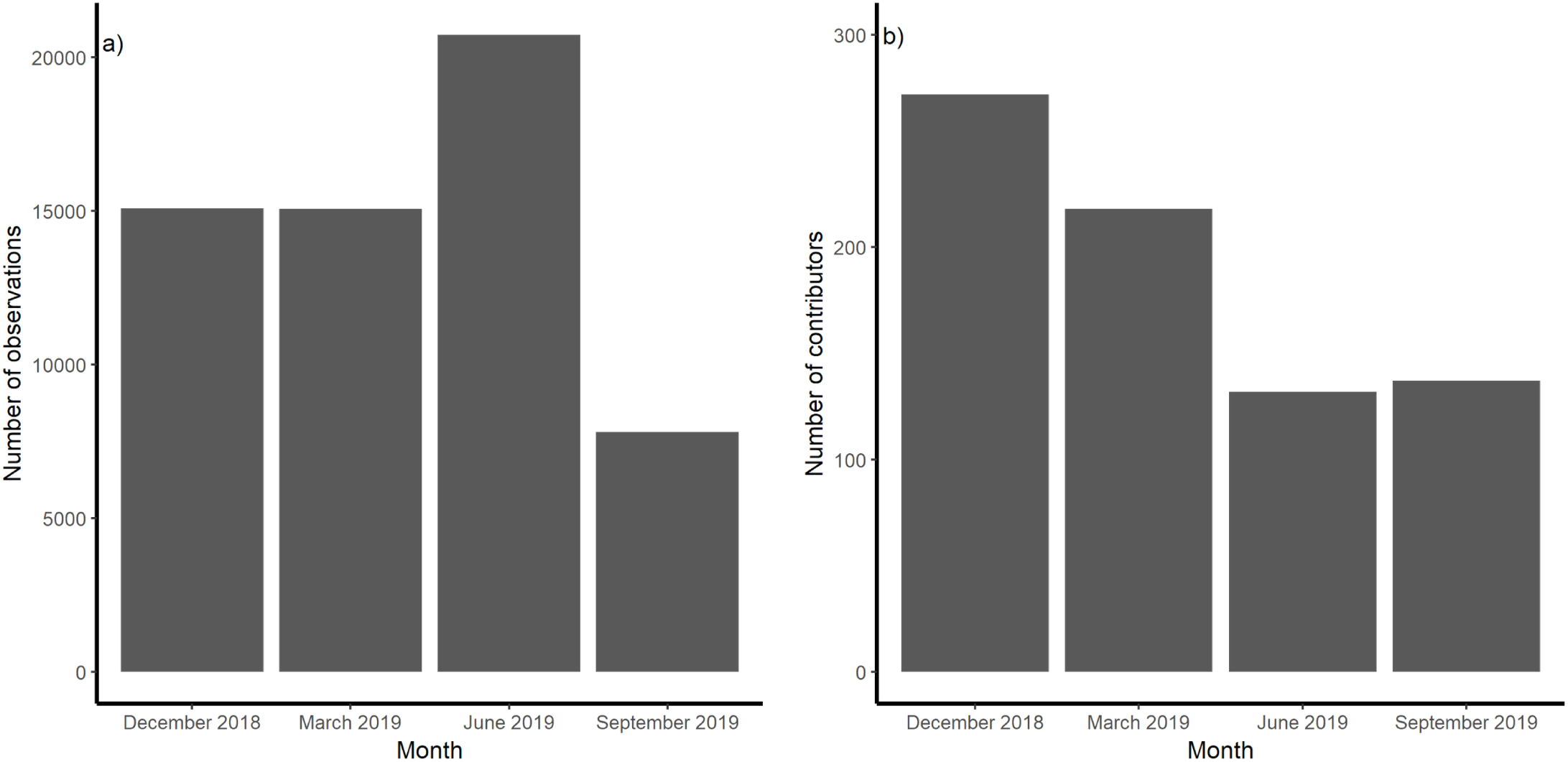
Number of observations (a) and number of participating contributors (b) in four bioblitz events held across India between December 2018 and September 2019.

*Mangifera indica* and *Tectona grandis* were among the top 10 most reported species in all four events, and we present results for only *M. indica* here. The fruiting phenology of *M. indica* was found to vary with latitude across 4 different months of assessment (Figure 6). In December 2018 and March 2019, a small proportion of trees were in fruit, and these were primarily in southern India. In June 2019, trees were in fruit across all latitudes and by September 2019, fruiting in *M. indica* had receded, with less than 2% of trees in fruit, again restricted to the southernmost latitudes.

**Figure 6:**
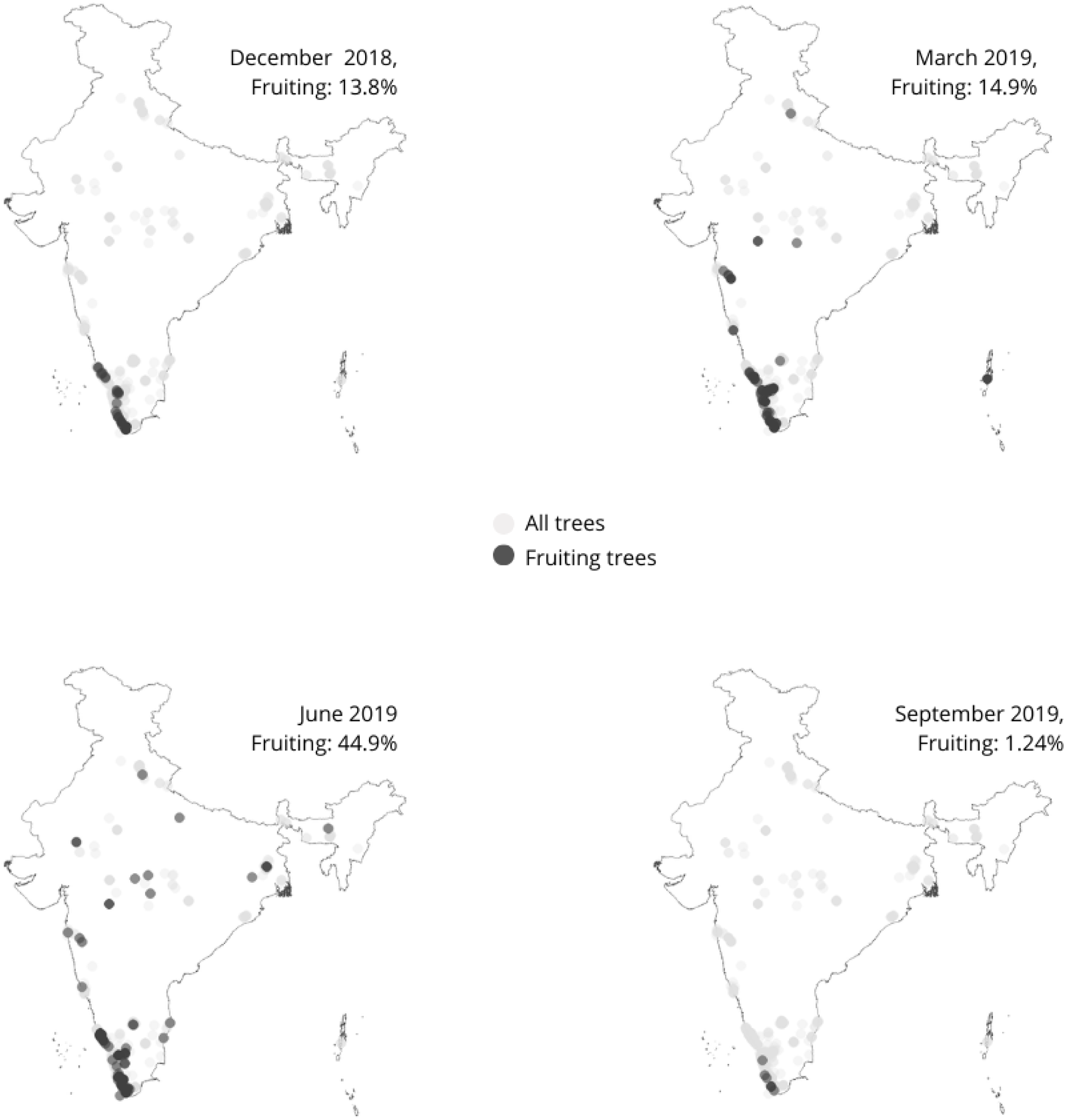
Spatial variation in the fruiting phenology of *Mangifera indica* (all varieties) as recorded during 4 bioblitz events in India. The south-western part of the country shows the presence of fruits on trees nearly all through the year, while fruiting is much more seasonal (between March and June) in central and northern India

## Discussion

Our first description of a long term, India-wide citizen science dataset contributes towards a basic understanding of seasonal phenology of common, widespread trees. Across temperate latitudes, historical information on plant phenology has been used as a reference to compare contemporary phenological changes and also identify climatic correlates of these changes. In the tropics, and especially in India, a similar baselines for species phenology across latitudes and seasons is lacking (Ramaswami et al 2019). The citizen science study presented here provides information on four widespread, common tropical species, which can potentially serve as references for future changes in phenology. For instance, in *A. heterophyllus* and *M. indica*, flowering and fruiting peaks in Kerala are well-defined over time, and can be used to detect shifts in peaks observed in the future.

Current phenological patterns can be compared against anecdotal time-points (such as plant phenology traditionally associated with festivals) in order to assess possible shifts in the reproductive phenology of culturally important trees. For instance, SeasonWatch data can be used to provide quantitative support to the anecdotal observations on changes in flowering dates of species such as *C. fistula*. In India, apart from Vishu, several seasonal festivals are associated with the appearance of flowers on culturally important tree species. The festival of Holi in north India is associated with flowering in *Butea monosperma* (Lam.) Taub. (Flame-of-the-forest tree); the flowers of which are collected and used to create a dye used in the festivities (Anonymous 2015). Similarly, the Kannada new year *Ugadi* and the Tamil harvest festival *Pongal* are associated with flowering in *Azadirachta indica* (Neem tree).

Latitudinal patterns in the vegetative and reproductive phenology of widespread and abundant species can help discern the range of underlying environmental factors affecting the appearance of phenophases. In temperate regions, there is typically a negative relationship between autumn phenology and latitude, with northern populations showing the appearance of phenophases such as leaf colour-change and leaf-fall before southern populations. For eg, in Japan, the timing of fall-leaf phenology was found to be negatively associated with latitude and temperature in the temperate species *Acer palmatum* and *Gingko biloba* over the duration of 53 years (Doi et al 2008).. In tropical regions, on the other hand, where spring and autumn temperature changes are not as stark as in temperate latitudes, vegetative phenology such as leaf-out may be triggered, instead, by photoperiod or preceding the onset of monsoons across different latitudes (Adole et al., 2019; Elliott et al., 2006; Ryan et al., 2017). In SeasonWatch, however, we were unable to detect any changes in vegetative phenology with latitude due to the large time-gaps between consecutive bioblitz assessments. Reproductive phenology is also associated with latitude in temperate regions, with some herbaceous species such as *Lythrum salicaria* exhibiting growth and flowering phenology earlier in northern latitudes (Olssen et al 2002). In other species, such as *Crataegus monogyna*, the opposite pattern has been reported, with southern populations showing fruiting phenology before northern populations (Guitian 1998). *M. indica*, shows the latter pattern, with southern populations showing reproductive phenophases before the northern populations, indicating a possible link with the Indian monsoon. In SeasonWatch, *M. indica* includes cultivated varieties as well, which may have variety-specific fruiting phenology.

The initial description of tropical tree phenology merits further investigation in terms of underlying environmental correlates, such as temperature, precipitation, and photoperiod. Given the large spatial skew in SeasonWatch data, correlational inferences at the scale of the country are likely to be affected by high variability. There is a better scope, however, for exploring temporal patterns of environmental changes on tree phenology, at least from the state of Kerala, with the addition of a few more years of data. A future research question could be addressing the changes in tree phenology in relation to changing climate. Overall changes in tropical tree phenology are expected to cascade through trophic levels. At present the effects of phenological change in tropical tree species from the Indian subcontinent on the life cycles of animals is poorly known. A research question that emerges thus, is, what are the downstream trophic impacts of tree phenology and what changes are likely to occur in these interactions given climate change.

## Conclusion

Citizen science has the potential to contribute immensely to mainstream scientific understanding of tropical tree phenology via building baselines. However, a citizen science project such as SeasonWatch faces several challenges, including - ensuring sufficient sample sizes per species per season from different locations, as well as data quality and accuracy. Managing these challenges is especially difficult in the Indian context given the high diversity of languages and the unequal access to the internet, as well as mass and social media, and at times literacy. It is therefore not possible to sustain such programmes without initiating and maintaining long-term partnerships with regional groups and organisations. Despite increasing outreach in multiple languages to promote awareness about the project, providing technological support free of cost for contributors, and simple visual design of the user interface created by taking inputs from contributors during the testing stage, data quantity remains low and skewed to a single state in the country. Furthermore, mechanisms to ensure data quality need to be implemented at multiple stages of data contribution. With this paper, we extend a call for action to sustain long-term interest and participation to develop a baseline for common tropical tree species that can be used to understand long-term consequences of climate change on tropical tree phenology.

## Acknowledgements

We thank the numerous citizen scientists from across India and especially Kerala who have tirelessly contributed data to the SeasonWatch project, making this description possible. Project funding for the duration of the study came from Wipro Foundation. Muhammad Nizar, regional coordinator for Kerala and regional partners Mathrubhumi have ensured that citizen scientists engage with and contribute to the programme regularly. Veena HT and Farheen Anjum have created the SeasonWatch website and Android app that contributors use to upload data.

## Appendix 1

Number of trees observed per species per week varies throughout the year. The patterns reported for species in Fig 3 are thus based on varying sample sizes per week. Lowest number of trees observed are usually between weeks 10 and 30 of the year - coinciding with summer holidays in schools in southern India. Sample sizes are as follows for these species; note that Y axis values vary - a) *Mangifera indica*, b) *Artocarpus heterophyllus*, c) *Tamarindus indicus* and d) *Phyllanthus emblica*. Note that thought the overall sample sizes for all species seem to fluctuate nearly exactly, total trees observed (Y axis) differs with species, with most number of trees observed for *A. heterophylla* and the least for *P. emblica*

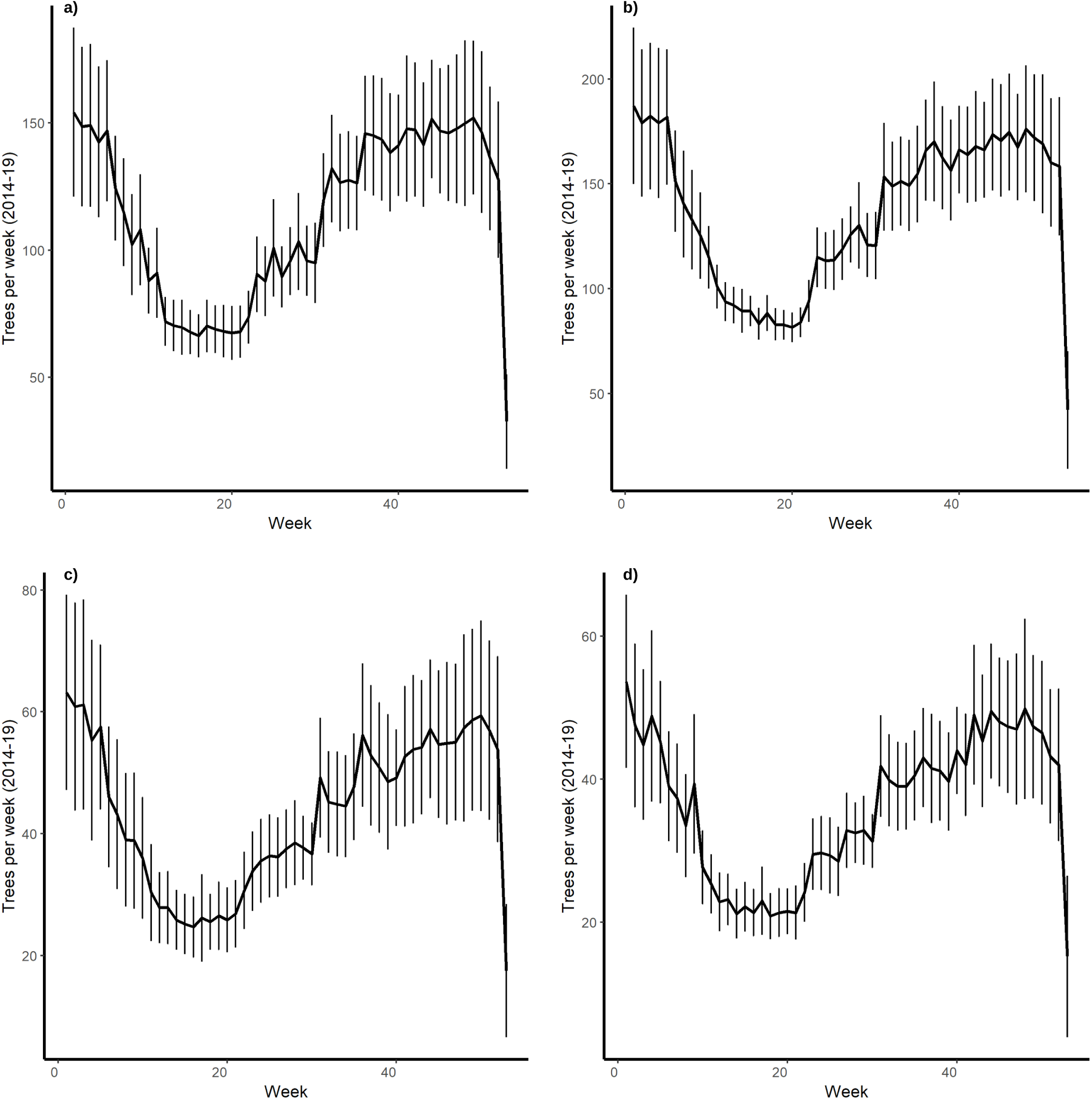

